# Newly identified SpoVAF/FigP complex: the role in *Bacillus subtilis* spore germination at moderate high pressure and influencing factors

**DOI:** 10.1101/2024.10.21.619482

**Authors:** Fengzhi Lyu, Ziqi Gong, Tianyu Zhang, Lei Rao, Xiaojun Liao

## Abstract

The SpoVAF/FigP complex, a newly identified ion channel, has been shown to amplify the response of germinant receptors (GRs) to nutrient germinants. However, its contribution to high-pressure-induced germination remains unexplored. In this study, we discovered that the 5AF/FigP complex plays an important role in the GRs-dependent germination of *Bacillus subtilis* spores under moderate high pressure (MHP) by facilitating the release of ions such as potassium (K^+^), a mechanism in parallel with its role in nutrient-induced germination. Despite its predicted function as an ion channel, the 5AF/FigP complex fails to respond to MHP in the absence of GerA-type GRs. We quantitatively examined the factors that influence the 5AF/FigP complex’s function in MHP-induced germination using modeling and fitting techniques. Our results indicate that the complex’s amplification effect is both enhanced and accelerated as pressure levels increase from 50 to 200 MPa. However, raising the MHP treatment temperature from 22 to 30°C only speeds up the complex’s function without enhancing its effectiveness. Moreover, extreme conditions of higher pressure (300 MPa) and temperature (34-37°C) can diminish the complex’s functionality. Additionally, the amplification effect is weakened in spores produced at both elevated and reduced sporulation temperatures. Taken together, our findings highlight the essential role of the 5AF/FigP complex in boosting the efficiency of MHP-induced germination. This revelation has enriched our understanding of the intricate mechanisms underlying GRs-dependent germination in *Bacillus* spores, offering valuable insights that can be utilized to refine the germination-inactivation strategies within the food industry.

## Introduction

Controlling bacterial spores presents a persistent challenge for food industry, as these dormant forms can survive most adversity stresses during food processing [1,2,3]. To address this concern, the food industry has adopted the germination-inactivation approach, which involves first inducing spores to germinate and then inactivating them through gentle processing techniques [4,5]. This approach is founded on the principle that once spores have fully germinated, they lose their resilience and become more vulnerable to milder inactivation treatments [6,7]. In this circumstance, achieving complete germination is essential for effective spore inactivation, and the efficiency of germination is crucial for the success of the germination-inactivation strategy [8].

The underlying mechanism of spore germination have been studied for decades [7,8,9,10]. Using *Bacillus subtilis* as a model, the germination process begins with the binding of nutrients to germination receptors (GRs), such as GerA, which specifically recognizes L-alanine as a germinant [11,12]. This triggers a series of events, including the activation of GRs, the release of pyridine-2, 6-dicarboxylic acid (DPA) in complex with Ca^2+^ (Ca-DPA), the hydrolysis of the spore cortex, and the rehydration of the spore core [7,8]. The final event marks the completion of germination and coincides with the loss of the spore’s resistance [7,8,13]. In addition to nutrients, physical stimuli such as high hydrostatic pressure (HHP) can also trigger spore germination [14,15,16]. The effects of HHP on spore germination differ depending on the pressure range: (i) at 50−300 MPa (moderate high pressure, MHP), it activates GerA-type GRs, similar to the mechanism of nutrient-induced germination with L-alanine [15,17,18]; (ii) at 400−600 MPa (very high pressure, VHP), it is supposed to open the SpoVA channel, releasing Ca-DPA and initiating germination [18]. As a commercial pasteurization method, HHP can effectively eliminate vegetative cells and germinated spores, although it has a limited effect on dormant spores [5,19]. Therefore, HHP is being actively explored for its potential role in the germination-inactivation strategy, as it has the dual capability of inducing germination and eliminating germinated spores [5,20,21]. Recently, Gao et al. (2024) have identified a novel germination component, the SpoVAF/FigP complex, which is composed of SpoVAF (5AF) and its indispensable partner protein FigP (YqhR) [22]. This complex functions as an ion channel during nutrient-induced germination, enhancing the process by facilitating the release of ions that amplify the response of germination receptors (GRs) to germinants [22]. It is hypothesized that the SpoVAF/FigP complex is activated by ions released through GerA-type GRs, thereby enhancing the germination signal. However, the role of the 5AF/FigP complex in germination induced by MHP is yet to be determined. Given the potential mechanistic differences in GRs activation between nutrient-induced and MHP- induced germination [8,15,17], it is not clear if this complex plays a similar role in both scenarios. Therefore, further research into the function of the 5AF/FigP complex during MHP-induced germination is crucial for gaining a comprehensive understanding of GRs-dependent germination mechanisms. Additionally, clarifying the roles of the 5AF/FigP complex could significantly contribute to the optimization of the germination-inactivation strategy employing HHP. This knowledge could lead to more effective methods for controlling spore germination and inactivation, ultimately enhancing food safety and quality.

In this study, we explored the function of the recently discovered 5AF/FigP complex in the germination of *Bacillus* spores under MHP, focusing on the release of Ca-DPA and potassium ions. We also explored various factors that could influence the activity of the 5AF/FigP complex during MHP-induced germination, considering the pressure level, pressurization temperature, and sporulation temperature. To further our analysis, we quantitatively modeled the DPA release curves, allowing us to assess the effects of these factors on germination rates and the overall percentage of germinated spores. This meticulous approach has yielded valuable insights that advance our understanding of the mechanisms behind HHP-induced germination in *Bacillus* spores. Our findings not only enrich the scientific community’s knowledge but also have practical implications for the food industry. By enhancing our grasp of the germination-inactivation strategy, we can improve the application of this method, leading to more effective spore control and ensuring greater food safety and quality.

## Results

### 5AF/FigP complex enhances MHP-induced germination efficiency likely via amplifying the germination responses of GerA-type GRs

In the 5AF/FigP complex, FigP was evidenced by Gao et al. (2024) as an essential partner protein of 5AF, with mutual dependence for co-localization and function [22]. Mutant spores lacking either the *spoVAF* (*5AF*) or *figP* gene exhibited similar germination defects as double mutant spores under L-alanine induction, indicating that the absence of either component protein resulted in the loss of function of the complex [22]. Coincidentally, FigP (YqhR) was also identified in our previous TMT proteomics analysis as a significant protein highly expressed during sporulation and affected spore germination [26]. Therefore, we decided to focus on FigP effect on MHP-induced germination. We initially constructed the Δ*figP* mutant and its complementation strains to examine their spore germination phenotypes under MHP treatments (200 MPa/26°C). The total DPA contents of both mutant spores were not significantly different (p > 0.05) from that of the wild-type (Fig. S1). Phase-contrast images of the Δ*figP* mutants revealed a notable decrease (p < 0.05) in the percentage of phase-dark spores after MHP treatment for 7 min compared to the wild-type (Figs. 1A-1B). Specifically, only 59.33% of the Δ*figP* mutant spores transformed to phase-dark, whereas the data of wild-type was 92.91% (Fig. 1B). Additionally, as shown in Fig. 1C, a remarkable decrease (p < 0.05) was observed in DPA release from the Δ*figP* mutant spores compared to the wild-type after MHP treatment for 10 min, with percentages of 29.5% and 63.5%, respectively. Importantly, complementation of *figP* in the Δ*figP* mutant rescued the germination defect of according spores to that of the wild-type (Figs. 1A-1C). These findings proved that the absence of *figP* in spores resulted in significant germination defects under MHP treatments, indicating the crucial role of the 5AF/FigP complex in MHP-induced germination.

**Figure 1.**
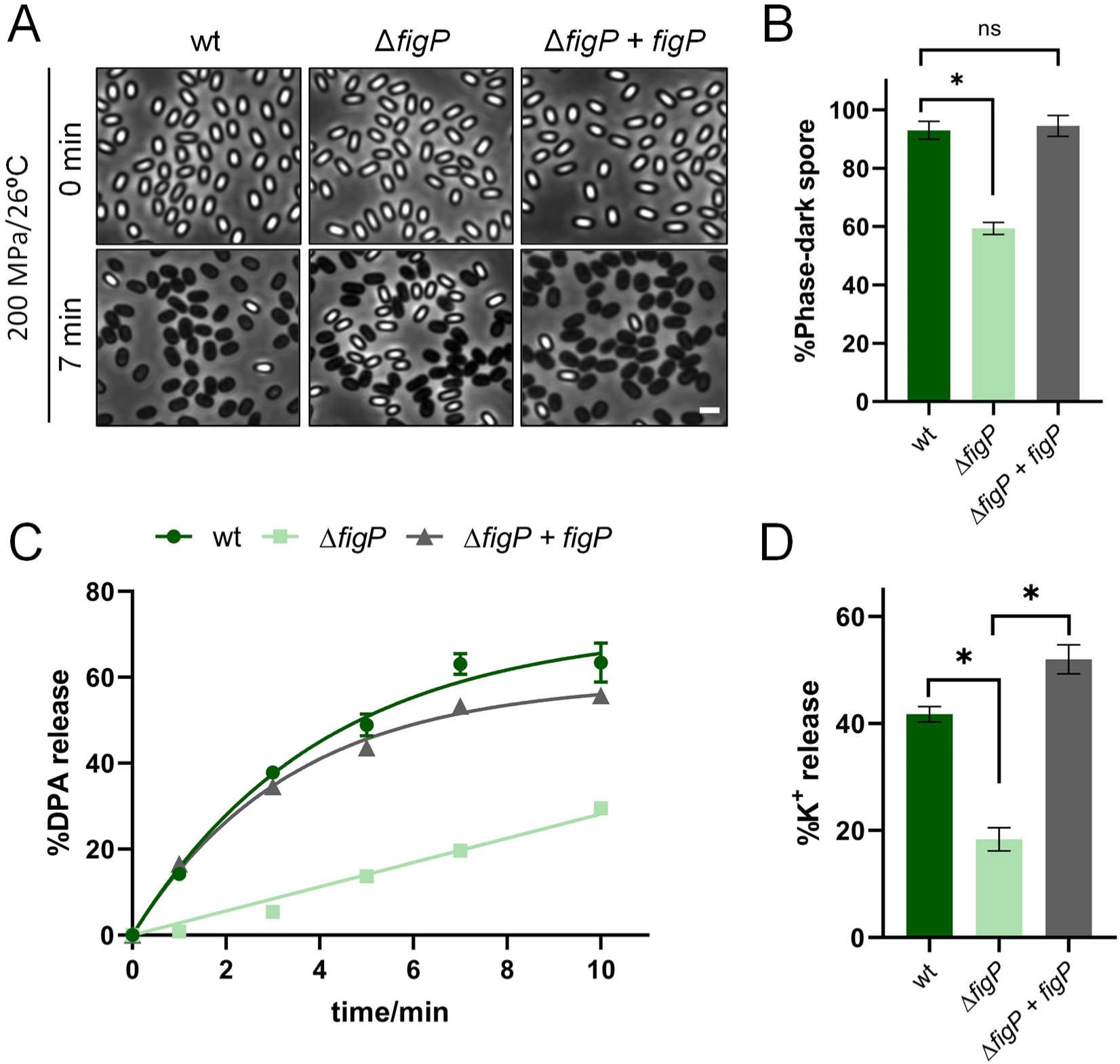
The function of SpoVAF/FigP complex in *B. subtilis* spore germination at MHP. (A) Representative phase-contrast images of *B. subtilis* PY79 (wt), YZ03 (Δ*figP*), and YZ05 (Δ*figP, amyE::figP*) spores before and after 7-min MHP treatment (200 MPa/26°C). Scale bar, 2 μm. (B) Quantification of phase-dark spores shown in (A). Data are presented as percentages of the number of the phase-dark spores and all spores in the same image (n ≥ 800 for each strain). (C) The percentage DPA release at 0, 1, 3, 5, 7, 10-min MHP treatment, relative to the total DPA content in spores described in (A). See also Figure S1. (D) The percentage K^+^ release at 1-min MHP treatment, relative to the total K^+^ content in spores described in (A). See also Figure S2. For (B-D), data are represented as mean ± SD. Shown are the representative results of three independent biological experiments, each with three replicates. One-way ANOVA with Tukey’s multiple comparisons test was performed to compare the significant differences. Asterisks denote the significance levels: *p < 0.05.

In nutrient-induced germination, the 5AF/FigP complex is identified as an ion channel that releases monovalent cations such as K^+^ to enhance the response of GerA-type GRs to nutrient germinants, thereby elevating germination efficiency [22]. To verify whether this complex serves a similar role in MHP-induced germination, we analyzed the germination exudates from the mutant spores collected before and after MHP treatments. It was found that the release of K^+^ in the Δ*figP* mutant spores was significantly delayed compared to that in the wild-type, with percentages of 18.34% and 41.71%, respectively, after 1-min MHP treatment (Fig. 1D). Moreover, to validate our findings, we examined the MHP-induced germination phenotype of the Δ*5AF* mutant spores. Interestingly, both DPA and K^+^ release in the Δ*5AF* mutants resembled those in the Δ*figP* mutant under MHP treatment (Fig. S2). These results indicated that the disruption of the 5AF/FigP complex in spores led to delayed ion release such as K^+^ under MHP treatment. This suggested the potential role of the complex as mediating ion release during MHP-induced germination, which resembled its function in nutrient-induced germination. Therefore, ion release mediated by this complex was crucial in both nutrient- and MHP-induced germination to potentially amplify GerA-type GRs’ responses to germination stimuli.

To further verified the potential amplification effect of 5AF/FigP complex in MHP-induced germination, spores lacking all GerA-type GRs (Δ*gerBB* Δ*gerKB* Δ*yfkT* Δ*yndE* Δ*gerA*, Δ5) and spores lacking both GerA-type GRs and FigP (Δ5 Δ*figP*) were employed to complement with *gerA* from *B. subtilis* and *B. cereus*. Absence of FigP in these complemented spores had no effect on their DPA contents (Fig. S3). As shown in Fig. 2, the Δ5 mutant harboring *gerA* from *B. subtilis* exhibited greater germination efficiency in the presence of FigP compared to its absence under MHP treatment, with nearly one-fold increase in DPA release than the latter. Similar results were observed in the germination of the Δ5 and Δ5 Δ*figP* mutants complemented with *gerA* from *B. cereus*. These findings suggested that the amplification effect of the 5AF/FigP complex significantly enhanced the response of both GerA-type GRs to MHP induction. Taken together, the 5AF/FigP complex is essential for GRs-dependent germination of *Bacillus* spores under MHP, which regulates ion release to facilitate overall germination efficiency.

**Figure 2.**
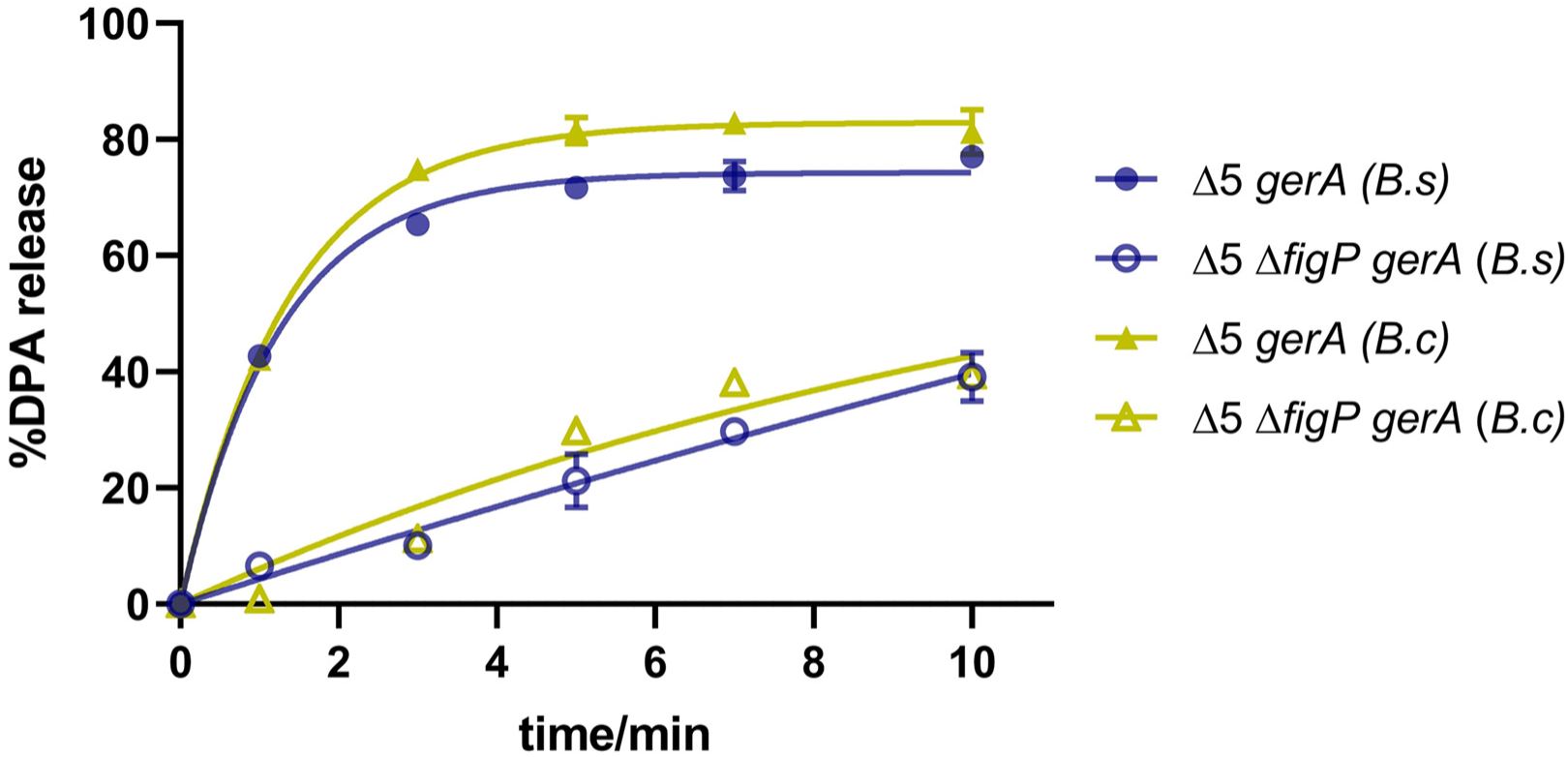
SpoVAF/FigP complex enhances MHP-induced germination of *B. subtilis* spores with a nonnative GerA. The percentage DPA release at 0, 1, 3, 5, 7, 10-min MHP treatment (200 MPa/26°C), relative to the total DPA content in spores of TYZ12 (Δ5, *amyE::gerA from B. subtilis*), YZR34 (Δ5 Δ*figP*, *amyE::gerA from B. subtilis*), TYZ13 (Δ5, *amyE::gerA from B. cereus*), and YZR37 (Δ5 Δ*figP*, *amyE::gerA from B. cereus*). See also Figure S3. Data are represented as mean ± SD. Shown are the representative results of three independent biological experiments, each with three replicates.

### 5AF/FigP complex is incapable of responding to HHP in the absence of GerA-type GRs

According to Gao et al. (2024), 5AF protein shared homology with A subunit of GerA, and formed ion channel region of the 5AF/FigP complex [22,27]. Since MHP can activate GerA-type GRs [15,28], we wonder whether the 5AF/FigP complex can be independently activated by HHP to induce germination. If this was the case, the absence of 5AF/FigP complex should cause HHP germination defect without GRs. Here, Δ5 and Δ5 Δ*figP* spores, possessing similar DPA content (Fig. S4), were employed to investigate the 5AF/FigP complex response to HHP without the influence of GerA-type GRs. As shown in Fig. 3, partial DPA was released for all mutant spores treated at 100-300 MPa over 10 min. Specifically, Δ5 spores released 2.1%, 8.7%, and 31.1% DPA, at 100, 200, and 300 MPa, respectively. Comparatively, the DPA release of Δ5 Δ*figP* spores was slightly increased by 2-4% (Fig. 3, Table 1). When the pressure elevated to over 400 MPa, the DPA release of Δ5 spores increased to 55.7-78.6%. Remarkably, the lacking of FigP in Δ5 spores resulted in 14-15% increase of DPA release. However, the lacking of 5AF in Δ5 spores showed no effect on both 200 and 500 MPa induced germination (Fig. S5). These results indicated that the 5AF/FigP complex could not independently respond to HHP in GRs-less spores, and the absence of FigP even somehow enhance the germination efficiency.

**Figure 3.**
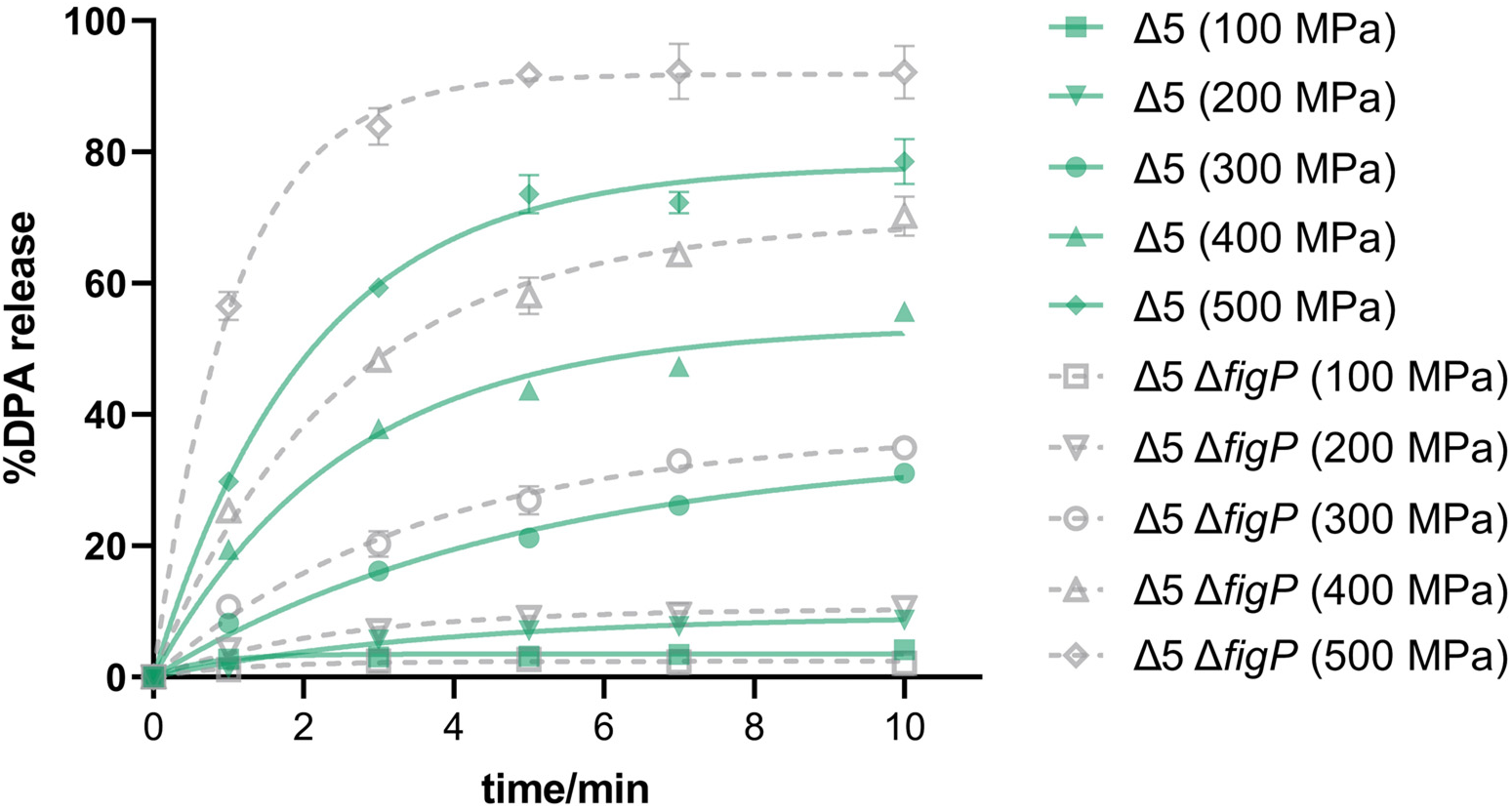
SpoVAF/FigP complex is incapable of responding to HHP in the absence of GerA. The percentage DPA release at 0, 1, 3, 5, 7, 10-min HHP treatment (100 - 500 MPa/26°C), relative to the total DPA content in spores of BLA201 (Δ5) and YZR33 (Δ5 Δ*figP*). See also Figure S4-S5. Data are represented as mean ± SD. Shown are the representative results of three independent biological experiments, each with three replicates.

**Table 1.** Kinetic constants and goodness of fit of DPA release curves of Δ5 and Δ5 Δ*figP* spores under 100-500 MPa, corresponding to Figure 3.

| Model |  | BoxLucas1 Model |  |  |  | Higuchi Model |  |  | Korsmeyer-Peppas Model |  |  |  | G<sub>max</sub> |
| --- | --- | --- | --- | --- | --- | --- | --- | --- | --- | --- | --- | --- | --- |
| | | $\frac{M_t}{M_\infty} = a(1 - e^{-bt})$ | | | | $\frac{M_t}{M_\infty} = kt^{0.5}$ | | | $\frac{M_t}{M_\infty} = kt^n$ | | | | |
| Pressure/MPa | Strains | *a* | *b* | Adj. R<sup>2</sup> | RMSE | *k* | Adj. R<sup>2</sup> | RMSE | *k* | *n* | Adj. R<sup>2</sup> | RMSE |  |
| 100 | Δ5 | 3.513 | 1.444 | 0.914 | 0.389 | 1.449 | 0.765 | 0.673 | 2.593 | 0.175 | 0.963 | 0.254 | 2.074 |
|  | Δ5 Δ*figP* | 2.395 | 0.818 | 0.394 | 1.036 | 0.921 | 0.307 | 1.162 | 1.613 | 0.190 | 0.342 | 1.080 | 4.180 |
| 200 | Δ5 | 9.406 | 0.264 | 0.972 | 0.552 | 2.878 | 0.937 | 0.863 | 2.638 | 0.547 | 0.933 | 0.847 | 8.701 |
|  | Δ5 Δ*figP* | 10.368 | 0.422 | 0.927 | 1.014 | 3.669 | 0.911 | 1.176 | 4.603 | 0.376 | 0.926 | 1.023 | 10.541 |
| 300 | Δ5 | 35.050 | 0.202 | 0.990 | 1.047 | 9.658 | 0.994 | 0.858 | 8.586 | 0.563 | 0.998 | 0.492 | 31.094 |
|  | Δ5 Δ*figP* | 37.400 | 0.275 | 0.987 | 1.404 | 11.684 | 0.984 | 1.653 | 11.942 | 0.488 | 0.982 | 1.645 | 34.937 |
| 400 | Δ5 | 53.439 | 0.394 | 0.985 | 2.295 | 18.620 | 0.976 | 3.045 | 22.250 | 0.403 | 0.988 | 2.031 | 55.705 |
|  | Δ5 Δ*figP* | 69.455 | 0.401 | 0.993 | 2.052 | 24.272 | 0.970 | 4.407 | 29.400 | 0.395 | 0.985 | 3.002 | 70.195 |
| 500 | Δ5 | 77.993 | 0.485 | 0.993 | 2.340 | 28.339 | 0.927 | 8.012 | 37.222 | 0.351 | 0.959 | 5.736 | 78.567 |
|  | Δ5 Δ*figP* | 91.865 | 0.929 | 0.995 | 2.377 | 36.250 | 0.776 | 16.594 | 62.932 | 0.194 | 0.972 | 5.603 | 92.131 |

### Factors influencing the amplification effect of 5AF/FigP on MHP-induced germination

Given the potential amplification effect of the 5AF/FigP complex in MHP-induced germination, it is significant to identify factors influencing the function of this complex. To analyze these factors quantitatively, DPA release curves were fitted to calculate the rate and efficiency of germination. Three models, including Box-Lucas1, Higuchi, and Korsmeyer-Peppas [29,30], were evaluated to screen the most suitable one that fitted the DPA release kinetics under MHP treatment. To ensure accuracy, data from Fig. 3 were used for the comparison of model fitting because they contained different patterns of HHP-induced germination behavior. Adjusted R-squared (Adj. R²) and root mean square error (RMSE) were utilized to compare model applicability, where higher Adj. R² and lower RMSE indicated better model fitting. When Adj. R^2^ < 0.90 or G_max_ < 15%, the model cannot fit the data well. In this case, experimental data rather than fitted data will be used for analysis. As for G_max_ > 15%, the BoxLucas1 model achieved an Adj. R^2^ greater than 0.985 and an RMSE less than 2.295, while the Higuchi and Korsmeyer-Peppas models showed minimum Adj. R^2^ values of 0.970 and 0.982, respectively, with maximum RMSE values of 4.407 and 3.002 (Table 1). Hence, the BoxLucas1 model was identified as the most suitable one for DPA release kinetics under MHP treatment, and will be employed in subsequent experiments to evaluate the effects of various factors on the function of 5AF/FigP. Specifically, G_max_ and *K*_max_, respectively representing germination percentage and rate as described in Materials and Methods, were used to assess the germination efficiency.

The factors including pressure level and treatment temperature of MHP, as well as sporulation temperature, were investigated for their effects on the amplification effect of 5AF/FigP. To this end, Δ5 *gerA* and Δ5 Δ*figP gerA* spores were employed, and Δ5 and Δ5 Δ*figP* spores were utilized as controls. The differences of G_max_ or *K*_max_ values between Δ5 *gerA* and Δ5 Δ*figP gerA* spores indicate the response of the 5AF/FigP complex to different factors.

#### (i) Pressure level

Pressure level is a critical factor for HHP-induced germination [17]. Thus, we investigated whether the function of the 5AF/FigP complex is affected by pressure levels. As shown in Figure 4 and corresponding model fitting data (Table 2), the G_max_ values for Δ5 and Δ5 Δ*figP* mutant spores remained below 10% across the pressure range of 50 to 200 MPa. Notably, both the G_max_ and *K*_max_ values for Δ5 Δ*figP gerA* spores increased with pressure from 50 MPa to 200 MPa. Given that the effect of GerA intensified with this increasing pressure, the impact of the 5AF/FigP complex was quantified by comparing the difference between Δ5 *gerA* and Δ5 Δ*figP gerA* spores. The germination percentage (G_max_) in Δ5 *gerA* spores increased by 10.7%, 43.4%, and 34.8% at 50, 100, and 200 MPa, respectively, compared to Δ5 Δ*figP gerA* spores. This indicates that the effect of the 5AF/FigP complex enhanced with increasing pressure from 50 to 100 MPa but slightly decreased at 200 MPa. Moreover, although the *K*_max_ values for both types of spores were similar at 50 MPa, the Δ5 *gerA* spores increased by 16.8% and 45.4% at 100 and 200 MPa, respectively, suggesting an accelerated amplification effect of the complex with increasing pressure. However, when the pressure was increased to 300 MPa, partial germination occurred in both Δ5 and Δ5 Δ*figP* spores, which was similar to Fig. 3. Notably, the increase of G_max_ and *K*_max_ values in Δ5 *gerA* spores, both measured at 300 MPa, were impaired (Table 2), suggesting that the amplification effect of the 5AF/FigP complex is likely reduced at this pressure. Taken together, the data indicated that the amplification effect of the 5AF/FigP complex can be enhanced and accelerated with pressure levels ranging from 50 to 200 MPa during MHP-induced germination. However, this effect was somehow impaired when pressure raised to 300 MPa.

**Figure 4.**
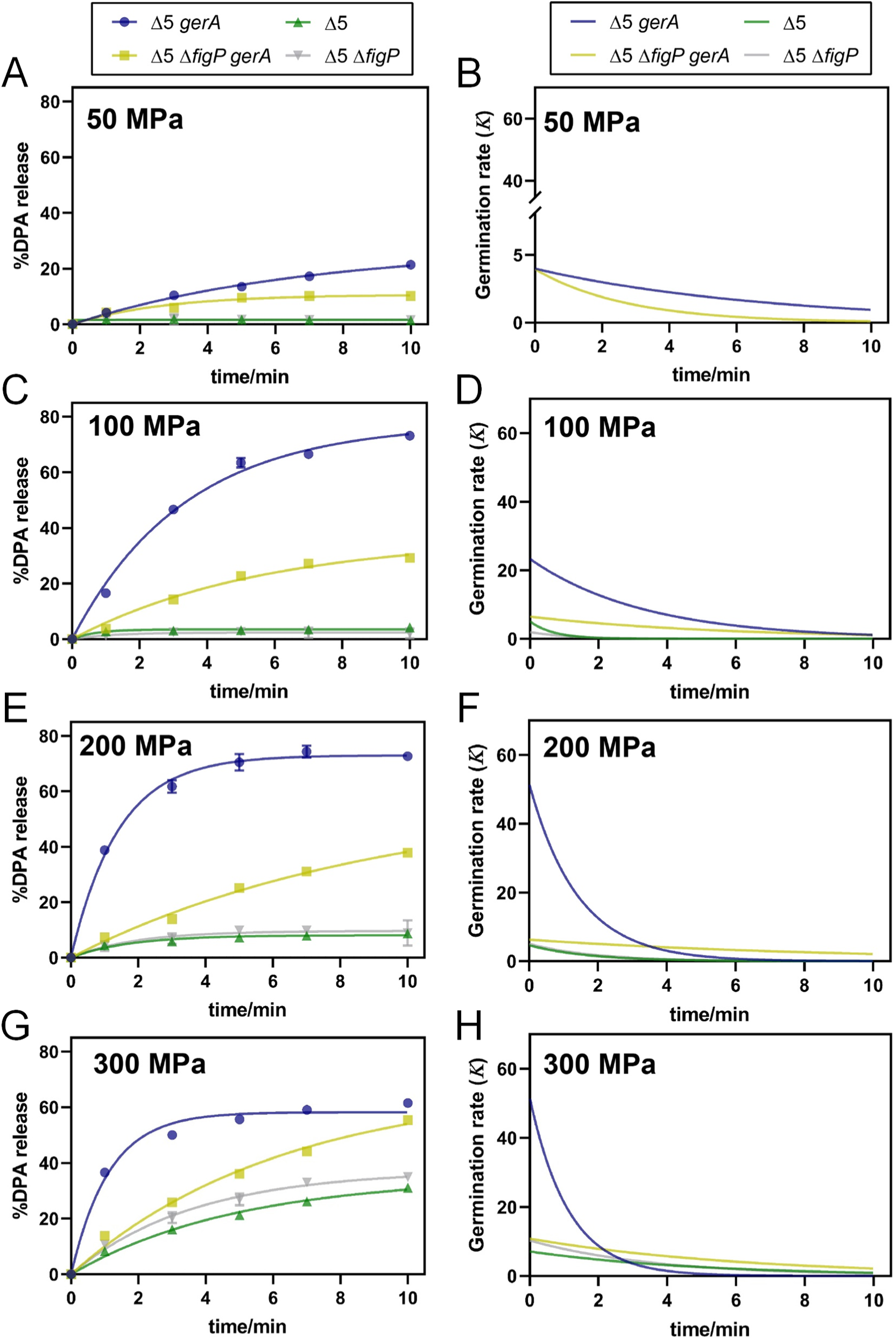
Effects of pressure level on the amplification effect of SpoVAF/FigP complex in MHP-induced germination. DPA release curves expressed as %DPA release were fit to BoxLucas1 model (A, C, E, and G) and calculated the derivative of the fitted curves to represent germination rate (*K*) at different time (B, D, F, and H). The percentage DPA release under 0, 1, 3, 5, 7, 10-min MHP treatment at 26°C, relative to the total DPA content in spores of TYZ12 (Δ5, *amyE::gerA*) and YZR34 (Δ5 Δ*figP*, *amyE::gerA*), was employed to fitted. Spores of BLA201 (Δ5) and YZR33 (Δ5 Δ*figP*) were used as controls and fitted by the same model. (A, B) 50 MPa; (C, D) 100 MPa; (E, F) 200 MPa; (G, H) 300 MPa. Data are represented as mean ± SD. Shown are the representative results of three independent biological experiments, each with three replicates.

**Table 2.** Maximum Germination efficiency (Gmax) and maximum germination rate (*K*max) of Δ5 *gerA* and Δ5 Δ*figP gerA* spores under 50-300 MPa at 26°C obtained after fitting germination curves to the BoxLucas1 model (Eq. 1), corresponding to Figure 4.

| Strains | Pressure/MPa | Constant | | Goodness of fit | | $K_{\max}$ | $G_{\max}$ |
| --- | --- | --- | --- | --- | --- | --- | --- |
| | | *a* | *b* | Adj. $R^2$ | RMSE | | |
| $\Delta 5$ *gerA* | 50 | 27.590 | 0.144 | 0.996 | 0.507 | $3.982 \pm 0.090^c$ | $21.072 \pm 0.292^c$ |
| | 100 | 77.587 | 0.300 | 0.993 | 2.275 | $23.283 \pm 0.542^b$ | $73.726 \pm 0.937^a$ |
| | 200 | 72.994 | 0.707 | 0.994 | 2.027 | $51.616 \pm 1.662^a$ | $72.936 \pm 0.155^a$ |
| | 300 | 58.205 | 0.888 | 0.984 | 2.643 | $51.712 \pm 0.341^a$ | $58.197 \pm 0.219^b$ |
| $\Delta 5$ $\Delta figP$ *gerA* | 50 | 10.655 | 0.371 | 0.958 | 0.765 | $3.948 \pm 0.165^C$ | $10.142 \pm 0.376^D$ |
| | 100 | 36.573 | 0.177 | 0.985 | 1.376 | $6.468 \pm 0.134^B$ | $30.335 \pm 0.250^C$ |
| | 200 | 57.628 | 0.109 | 0.992 | 1.203 | $6.258 \pm 0.117^B$ | $38.175 \pm 0.038^B$ |
| | 300 | 67.615 | 0.160 | 0.989 | 1.950 | $11.503 \pm 0.817^A$ | $52.029 \pm 2.285^A$ |

<sup>a-c</sup>For each row, means followed by the same letter are not significant different ( $p > 0.05$ ).
<sup>A-D</sup>For each row, means followed by the same letter are not significant different ( $p > 0.05$ ).

#### (ii) MHP Treatment temperature

In MHP-induced germination, treatment temperature is another critical factor influencing spore germination efficiency [17]. Thus, we investigated whether the function of 5AF/FigP complex under MHP treatment (200 MPa/10 min) could be affected by varying temperatures. As shown in Fig. 5 and corresponding model fitting data (Table 3), the Gmax values for Δ5 Δ*figP gerA* spores showed no significant differences (p > 0.05) across treatment temperatures of 22 to 34°C, ranging from 37.4% to 43.8% (Fig. 5, Table 3). Similarly, the *K*_max_ values exhibited minimal variation, with slight fluctuations between 5.5 and 8.8 over the same temperature range (Fig. 5, Table 3). These relatively stable germination parameters suggested that the function of GerA itself in MHP-induced germination was likely unaffected by treatment temperature ranging from 22 to 34°C. However, when the treatment temperature elevated to 37°C, both G_max_ and *K*_max_ values of these spores decreased significantly (p < 0.05). These reductions might be attributed to the negative effect of the higher temperature on GerA function.

**Figure 5.**
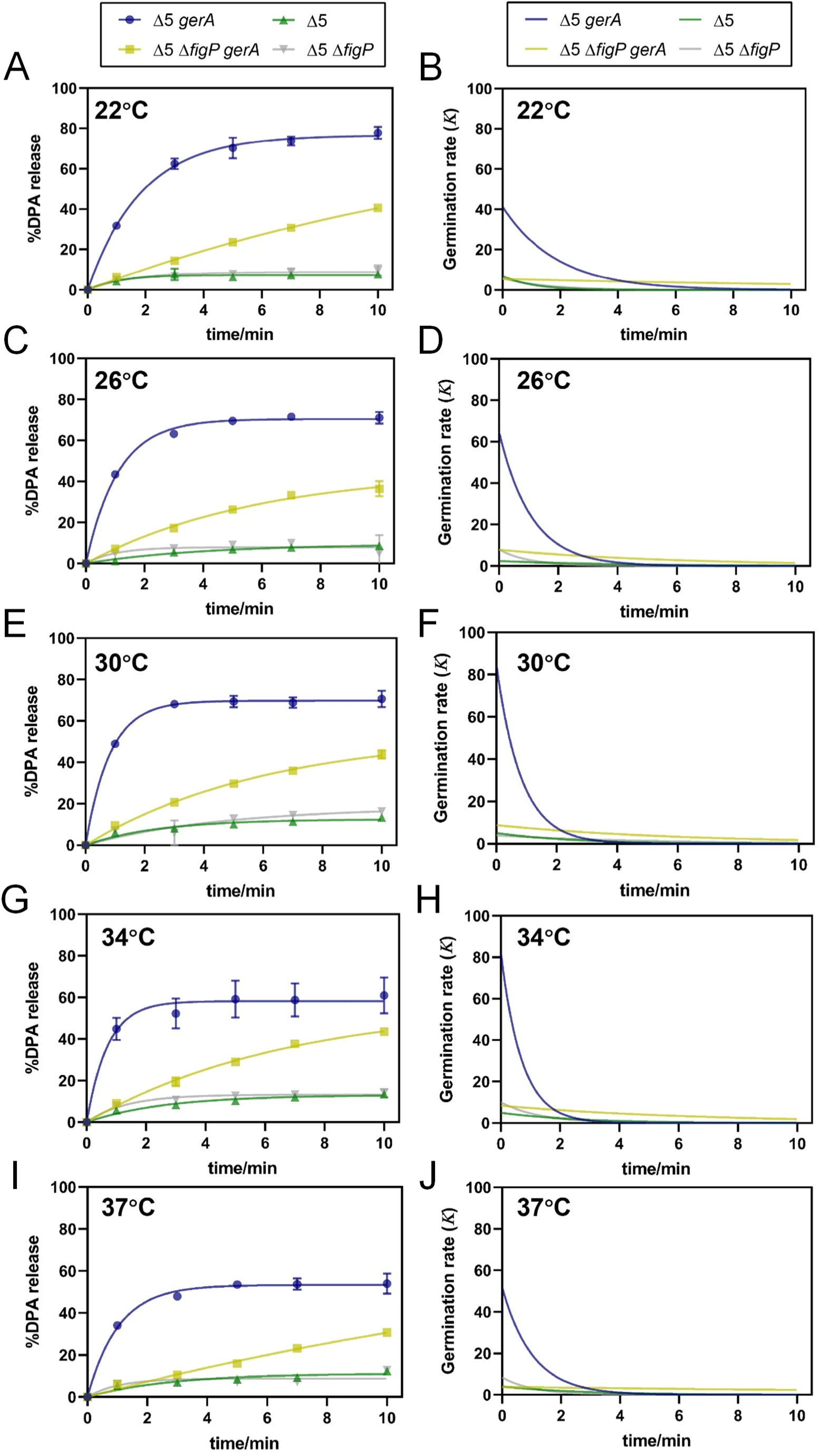
Effects of MHP treatment temperature on the amplification effect of SpoVAF/FigP complex in MHP-induced germination. DPA release curves expressed as %DPA release were fit to BoxLucas1 model (A, C, E, G, and I) and calculated the derivative of the fitted curves to represent germination rate (*K*) at different time (B, D, F, H, and J). The percentage DPA release under 0, 1, 3, 5, 7, 10-min 200 MPa treatment at varying temperature, relative to the total DPA content in spores of TYZ12 (Δ5, *amyE::gerA*) and YZR34 (Δ5 Δ*figP*, *amyE::gerA*), was employed to fitted. Spores of BLA201 (Δ5) and YZR33 (Δ5 Δ*figP*) were used as controls and fitted by the same model. (A, B) 22°C; (C, D) 26°C; (E, F) 30°C; (G, H) 34°C; (I, J) 37°C. Data are represented as mean ± SD. Shown are the representative results of three independent biological experiments, each with three replicates.

**Table 3.** Maximum Germination efficiency (Gmax) and maximum germination rate (*K*max) of Δ5 *gerA* and Δ5 Δ*figP gerA* spores under 200 MPa at 22-37°C obtained after fitting germination curves to the BoxLucas1 model (Eq. 1), corresponding to Figure 5.

| Strains | Temperature/°C | Constant | | Goodness of fit | | $K_{\max}$ | $G_{\max}$ |
| --- | --- | --- | --- | --- | --- | --- | --- |
| | | *a* | *b* | Adj. $R^2$ | RMSE | | |
| $\Delta 5$ *gerA* | 22 | 76.638 | 0.538 | 0.993 | 2.313 | 41.159 ± 5.366<sup>d</sup> | 76.463 ± 1.274<sup>a</sup> |
|  | 26 | 70.414 | 0.915 | 0.996 | 1.633 | 64.374 ± 1.264<sup>bc</sup> | 70.414 ± 0.971<sup>ab</sup> |
|  | 30 | 69.756 | 1.213 | 0.995 | 1.841 | 84.604 ± 0.363<sup>a</sup> | 69.761 ± 2.905<sup>abc</sup> |
|  | 34 | 58.220 | 1.410 | 0.930 | 5.781 | 82.158 ± 6.835<sup>ab</sup> | 58.223 ± 8.297<sup>bc</sup> |
|  | 37 | 53.388 | 0.968 | 0.988 | 2.099 | 51.758 ± 5.552<sup>cd</sup> | 53.454 ± 2.868<sup>c</sup> |
| $\Delta 5$ Δ*figP gerA* | 22 | 86.897 | 0.063 | 0.997 | 0.722 | 5.453 ± 0.017<sup>BC</sup> | 40.503 ± 1.150<sup>A</sup> |
|  | 26 | 45.507 | 0.171 | 0.989 | 1.408 | 7.822 ± 0.756<sup>AB</sup> | 37.354 ± 3.088<sup>AB</sup> |
|  | 30 | 53.748 | 0.163 | 0.994 | 1.164 | 8.783 ± 0.320<sup>A</sup> | 43.262 ± 1.788<sup>A</sup> |
|  | 34 | 56.489 | 0.149 | 0.993 | 1.287 | 8.424 ± 1.052<sup>A</sup> | 43.766 ± 0.024<sup>A</sup> |
|  | 37 | 83.329 | 0.046 | 0.987 | 1.175 | 3.823 ± 0.013<sup>C</sup> | 30.657 ± 0.745<sup>B</sup> |

<sup>a-d</sup>For each row, means followed by the same letter are not significant different ( $p > 0.05$ ).
<sup>A-C</sup>For each row, means followed by the same letter are not significant different ( $p > 0.05$ ).

Importantly, as the treatment temperature increased from 22 to 30°C, the *K*_max_ value of Δ5 *gerA* spores, which possessed both GerA and 5AF/FigP channels during MHP-induced germination, increased significantly (p < 0.05) from 41.2 to 84.6 (Fig. 5, Table 3), indicating a faster germination rate. Since GerA was insensitive to MHP treatment temperature within this range, it appeared that the increased germination rate was primarily attributed to the potentially enhanced function of 5AF/FigP complex. Specifically, the amplification effect of this complex was likely accelerated as temperature increased from 22 to 30°C. However, no significant difference (p > 0.05) was observed in the G_max_ value of Δ5 *gerA* spores within the same temperature range (Fig. 5, Table 3), which suggested that the percentage of germinated spores remained unchanged in the end of 10-min MHP treatment. Therefore, although the amplification effect of 5AF/FigP complex on germination rate might be improved by increasing MHP treatment temperatures from 22 to 30°C, its effect on the final germination ratio did not appear to be enhanced. However, once the treatment temperature exceeded 30°C, noticeable decreases were observed in both *K*_max_ and G_max_ value of Δ5 *gerA* spores, inferring impaired function of the 5AF/FigP complex at higher temperatures. Taken together, elevated treatment temperatures from 22 to 30°C could not impact the germination response of GerA to MHP, but likely accelerated the function of 5AF/FigP complex, leading to a faster germination rate. Nonetheless, this acceleration effect was likely weakened as temperatures continued to rise.

#### (iii) Sporulation temperature

Previous studies have reported that sporulation temperature impacts future spore’s germination capacity under MHP induction [28,31,32], which may impact the efficiency of germination-inactivation strategy mediated by MHP. To evaluated the effect of sporulation temperature on the function of the 5AF/FigP complex, Δ5 *gerA* and Δ5 Δ*figP gerA* mutants were induced to sporulate at 22, 37 and 44°C (referred to as S_22°C_, S_37°C_, and S_44°C_), and subsequently assessed for germination phenotypes under MHP induction (200 MPa/26°C/10 min). As illustrated in Fig. 6, both S_22°C_ spores exhibited remarkable germination defects, with G_max_ values of less than 11%. This indicated that the absence of *figP* in Δ5 *gerA* spores produced at a low temperature had no significant (p > 0.05) effect on their future germination under MHP induction (Table 4). Additionally, significant differences in germination efficiency were observed between two types of S_37°C_ spores under MHP treatment. Specifically, the *K*_max_ value of the Δ5 *gerA* mutant was calculated to be 10-fold higher than that of the Δ5 Δ*figP gerA* mutant during MHP-induced germination. Moreover, there was a nearly 30% increase in the G_max_ value in the Δ5 *gerA* mutant compared to the Δ5 Δ*figP gerA* mutant. These data suggested that the function of the 5AF/FigP complex during MHP-induced germination was greatly enhanced with increased sporulation temperature to 37°C. Interestingly, when the sporulation temperature was elevated to 44°C, the *K*_max_ value of the Δ5 *gerA* mutant was only 1-fold higher than that of the Δ5 Δ*figP gerA* mutant, whereas the lacking of *figP* in spores led to a slightly higher G_max_ value at the end of MHP treatment. The narrowed gaps in *K*_max_ values between the two types of S_44°C_ spores suggested a diminished role of the 5AF/FigP complex in MHP-induced germination. Taken together, these findings indicated that either lower or higher sporulation temperatures significantly compromised the function of the 5AF/FigP complex in MHP-induced germination.

**Figure 6.**
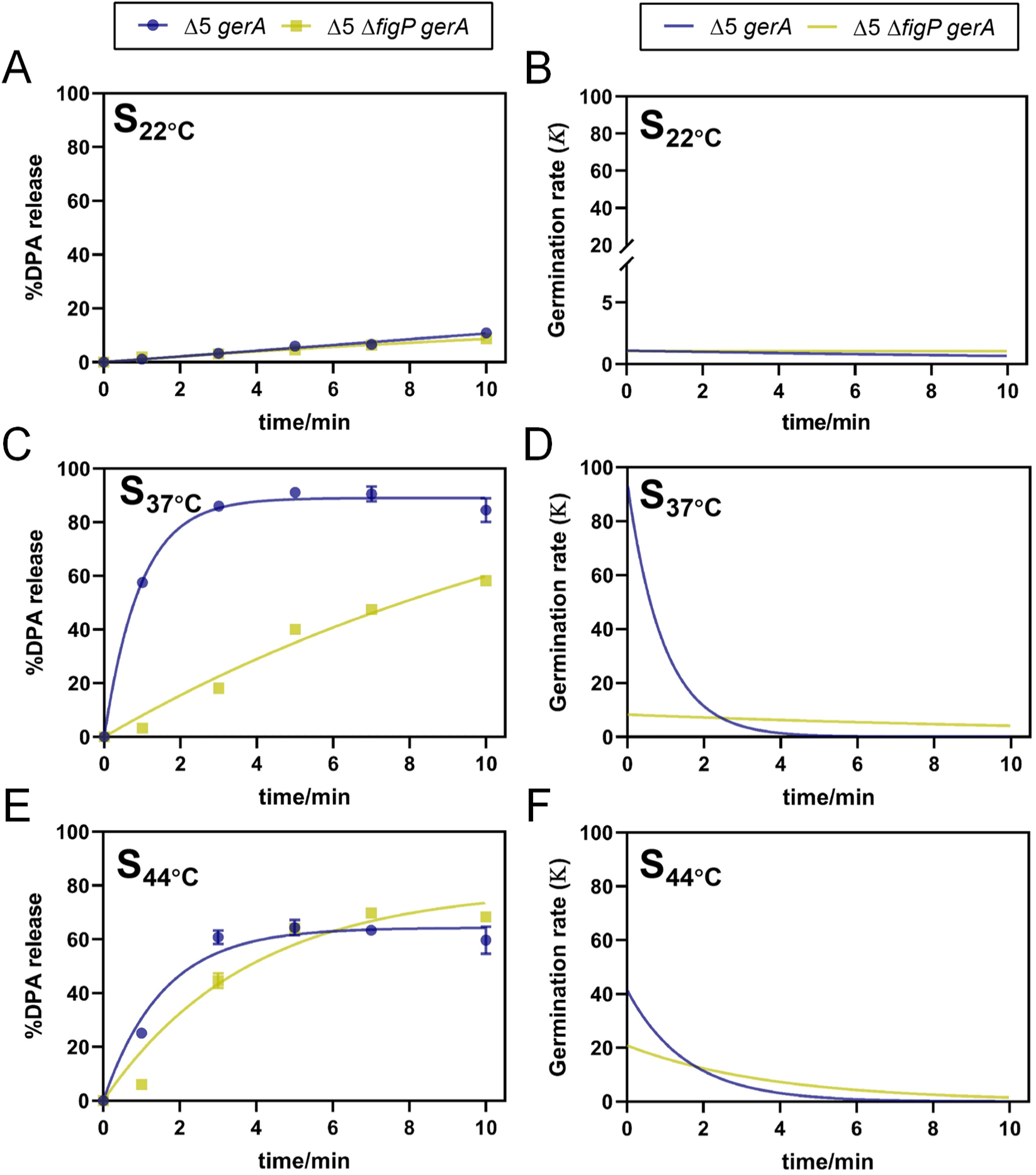
Effects of sporulation temperature on the amplification effect of SpoVAF/FigP complex in MHP-induced germination. DPA release curves expressed as %DPA release were fit to BoxLucas1 model (A, C, and E) and calculated the derivative of the fitted curves to represent germination rate (*K*) at different time (B, D, and F). The percentage DPA release under 0, 1, 3, 5, 7, 10-min 200 MPa treatment at 26°C, relative to the total DPA content in spores of TYZ12 (Δ5, *amyE::gerA*) and YZR34 (Δ5 Δ*figP*, *amyE::gerA*), was employed to fitted. (A, B) Sporulation temperature at 22°C; (C, D) Sporulation temperature at 37°C; (E, F) Sporulation temperature at 42°C. Data are represented as mean ± SD. Shown are the representative results of three independent biological experiments, each with three replicates.

**Table 4.**
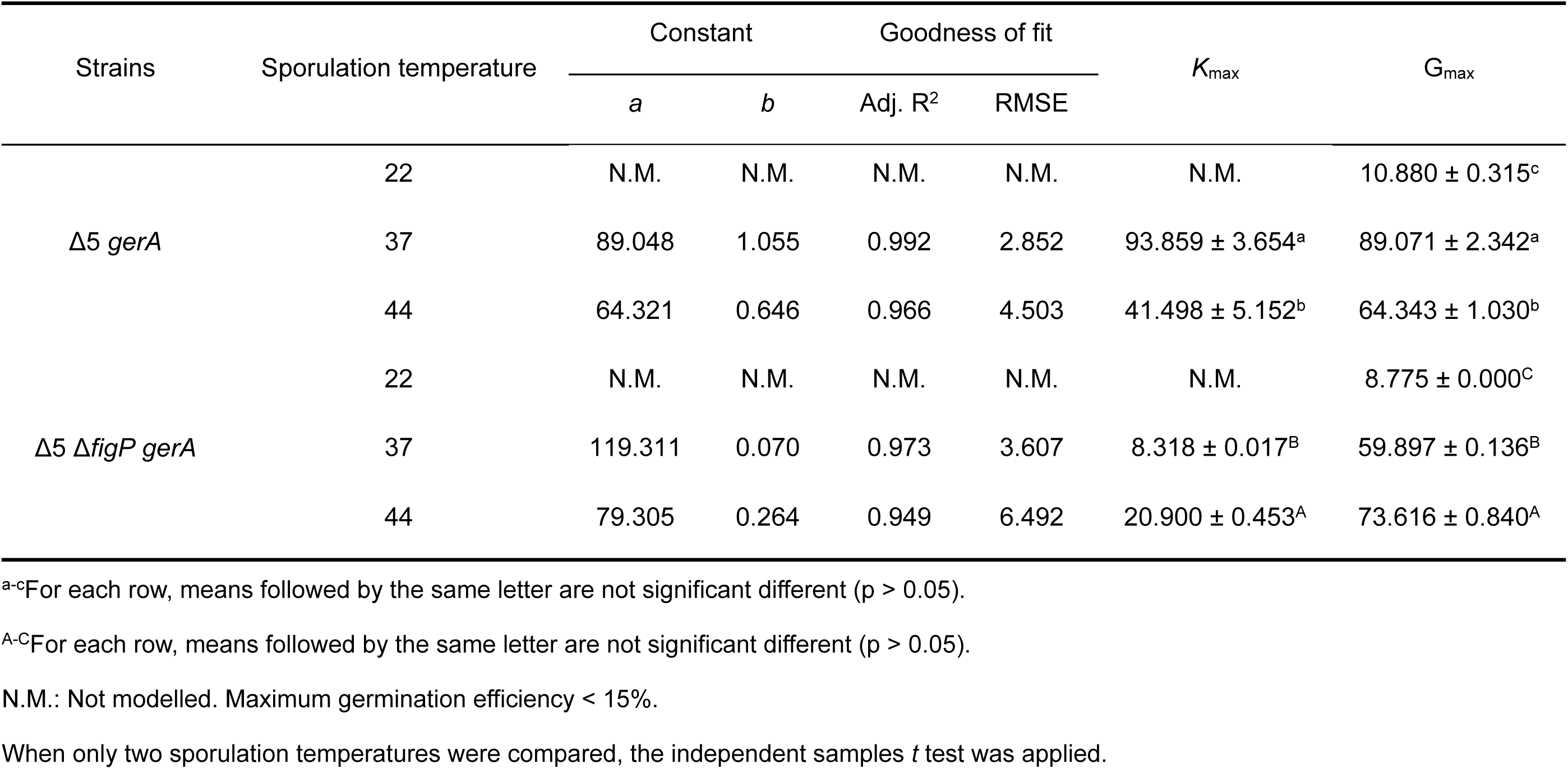
Maximum Germination efficiency (Gmax) and maximum germination rate (*K*max) of Δ5 *gerA* and Δ5 Δ*figP gerA* spores sporulated at 22-44°C under 200 MPa at 26°C obtained after fitting germination curves to the BoxLucas1 model (Eq. 1), corresponding to Figure 6.

## Discussion

In our study, we have revealed that the newly identified 5AF/FigP complex contributes to the germination of *B. subtilis* spores under MHP, mirroring its role in nutrient-induced germination (Fig. 7). While the 5AF/FigP complex is hypothesized to act as an ion channel, our findings indicate that it does not respond to MHP in the absence of GerA-type GRs. As show in Fig. 7, we observed that the complex’s amplification effect on germination is both enhanced and accelerated as the pressure levels rise from 50 to 200 MPa. However, increasing the MHP treatment temperature from 22 to 30°C only accelerates the function of the 5AF/FigP complex without enhancing its overall effectiveness. Notably, both higher pressures (300 MPa) and temperatures (34-37°C) were found to impair the complex’s functionality. Additionally, we discovered that the amplification effect of the 5AF/FigP complex can be suppressed in spores that are produced at either elevated or reduced sporulation temperatures. These findings provide valuable insights into the nuanced behavior of the 5AF/FigP complex and its role in the germination process under varying conditions.

**Figure 7.**
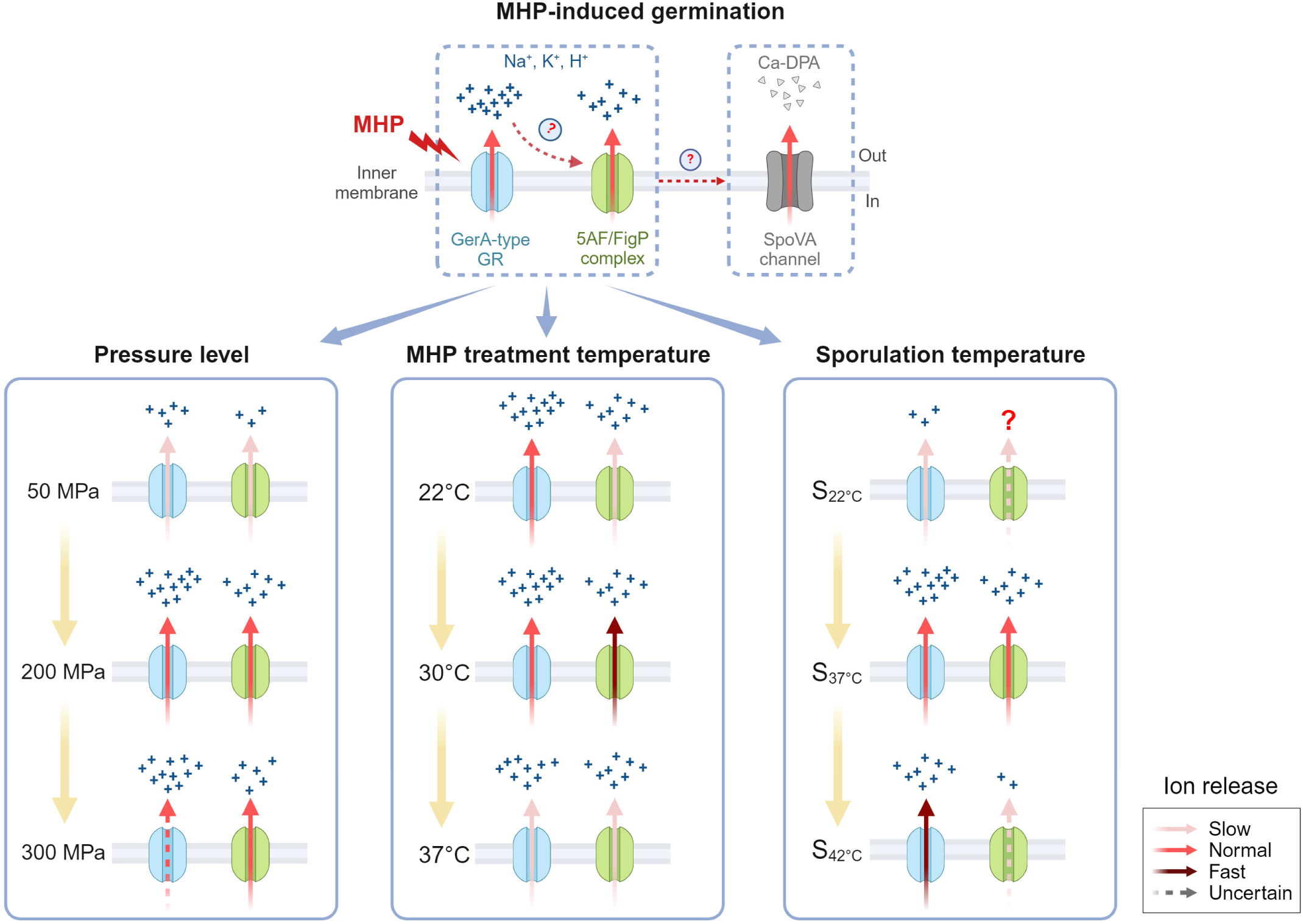
The model presenting the amplification effect of SpoVAF/FigP complex in MHP-induced germination, and the possible effects of pressure level, treatment temperature, and sporulation temperature on the function of the complex.

Interestingly, our research revealed that the absence of FigP paradoxically enhanced the germination efficiency of Δ5 spores, with this effect being particularly pronounced under VHP conditions (Fig. 3, Fig. S5, Table 1). Gao et al. (2024) have previously shown that 5AF and FigP are interdependent for stability [22], suggesting that the absence of FigP could lead to the defective assembly of the 5AF/FigP complex, thereby releasing 5AF. Our protein-protein interaction predictions using the STRING database (Fig. S7) [33], hint that 5AF might interact with other SpoVA subunits, particularly the channel-forming subunits C, D, and Eb [34], to modulate germination efficiency under VHP. However, the exact subcellular localization of the 5AF/FigP complex in relation to SpoVA channel subunits and the nature of their potential interactions await confirmation through experimental evidence. Moreover, the specific mechanism by which the SpoVA channel opens under VHP remains to be fully understood. Consequently, we currently lack a coherent model to explain the observed increase in DPA release under VHP in Δ5 spores lacking FigP, indicating a need for further investigation. This intriguing finding underscores the complexity of the germination process and the potential for novel insights into spore inactivation strategies.

Our experimental results demonstrated that 5AF/FigP complex was incapable of responding to HHP independently, whereas variations in MHP treatment conditions could influence the function of the complex. Specifically, increasing the pressure (from 50 to 200 MPa) both quickened and intensified the complex’s amplification effect. In contrast, elevating the MHP treatment temperature (from 22 to 30°C) solely augmented the rate of this effect. Gao et al. [22] predicted that the 5AF/FigP complex’s activation might rely on ions released from GerA-type GRs during nutrient-induced germination. This hypothesis elegantly accounts for our observations. As shown in Fig. 4 and Table 2, it is plausible that higher pressure levels expedite and enhance the release of ions from GerA, thus accelerating and strengthening the complex’s activation. However, an excessively high pressure (300 MPa), as reported by Heinz & Knorr [35], is not optimal for GerA activation, leading to a diminished activation efficiency and, by extension, a compromised function of the 5AF/FigP complex (Fig. 4, Table 2). In contrast to pressure, MHP treatment temperature exerts a minimal impact on GerA activation (Fig. 5, Table 3). Instead, it likely influences the 5AF/FigP complex’s function by affecting the velocity of ion movement and/or the complex’s intrinsic properties. As the temperature rises, the thermal motion of ions released from GerA may accelerate, potentially speeding the complex’s activation and, consequently, spore germination. An alternative explanation is that the 5AF/FigP complex could act as a temperature-sensitive ion channel. In this case, increased treatment temperatures might directly modify its protein conformation [36], to accelerate ion releasing efficiency. However, experimental evidence was needed to support these hypotheses.

Moreover, the sporulation temperature significantly influenced the 5AF/FigP complex’s performance during germination induced by MHP. As illustrated in Fig. 6 and Table 4, the complex in S_22°C_ spores exhibited minimal functionality, potentially due to the low activation level of GRs. In contrast, the function of the complex in S_44°C_ spores appeared to be somehow compensated, thereby its absence in Δ5 *gerA* spores did not significantly affect germination. Our findings could be attributed to the alterations in spore characteristics at different sporulation temperatures, encompassing: (i) fatty acid (FA) composition, where the anteiso-to-iso ratio and the level of unsaturated FAs in spore IM negatively correlated with sporulation temperature [37]; (ii) core DPA and water content, as DPA concentration increased while water content decreased with higher sporulation temperatures [38,39,40]; and (iii) spore coat structure, with lower sporulation temperatures promoting a more compact and less adherent outer coat, whereas higher sporulation temperatures leading to less abundant coat proteins [41]. However, due to the limited understanding of the activation mechanisms underlying MHP-induced germination, it remains challenging to predict how these spore property changes, induced by varying sporulation temperatures, precisely influence the functionality of the 5AF/FigP complex. Further research is needed to elucidate these complex interactions.

## Materials and methods

### Strains and plasmids

*B. subtilis* strains used in this study are listed in Table S1. Plasmids construction is described in Table S2, and primers are listed in Table S3. For gene replacement strategy, primer pairs were used to amplify the flanking genomic regions of the corresponding gene [23]. PCR products and the respective antibiotic resistance gene were used for Gibson assembly (NEB, USA) [24]. The product was used to transform *B. subtilis* PY79 to obtain the mutant allele.

### Spore preparation

Spores used in this study were prepared as described previously with some modifications [25]. Cultures of wild-type and mutant strains were cultivated in Luria-Bertani (LB) medium (Difco) at 37°C. Subsequently, cells (OD_600_ = 0.05) were inoculated into 100 mL flasks containing 15 mL Schaeffer’s liquid medium (Difco Sporulation Medium, DSM). Sporulating cultures were incubated at 37°C with shaking for 24 hrs. For study on the influence of sporulation temperature, sporulating cultures were incubated at 22°C, 37°C, and 44°C, respectively. Spores were harvested by centrifugation at 10,000 rpm for 10 min at 4°C and washed 3 times by sterile double distilled water (ddH_2_O). Then, spore purification was carried out followed Rao et al. [23]. Obtained spores were resuspended in ddH_2_O and kept in 4°C. After 7 days of washing, spores were centrifuged and purified by buoyant density centrifugation using Nycodenz. Briefly, the pellets were resuspended in 20% Nycodenz solution. Aliquots of the resuspension mixture (200 μL) were layered on top of 900 μL 50% Nycodenz in 1.5 mL centrifuge tubes, and tubes were centrifuged at 15,000 rpm for 20 min at 4°C. Pellets were collected and followed by at least 5 times washing using ddH_2_O. Spore purity was verified by phase-contrast microscopy. High purity spores (phase-bright spores > 99%) can be used in the following experiments, otherwise the purification process should be repeated.

### Spore germination induced by moderate high pressure (MHP)

Spores (OD_600_ = 0.5) were re-suspended with ddH_2_O and sealed in flexible plastic bags. The sealing bags containing spore suspension were placed in a high-pressure vessel with a volume of 5.0 L (Experimental High Pressure Equipment, SHHP-5L, Shanxi Leadflow Technology Co., Ltd, Shanxi, China), and subjected to pressure of 50 - 300 MPa for 1-10 min at 22-37°C as indicated in each experiment. The treated suspensions were placed on ice before centrifugation in tubes. Spore germination induced by MHP was detected by measuring the percentage of released DPA from the samples treated for 1, 3, 5, 7, and 10 min. The DPA release percentage was represented as the ratio of released DPA content at each time point to total DPA content.

### DPA measurement

DPA release was detected as described previously with some modifications [16]. Briefly, released DPA bound to Tb^3+^ and intense fluorescence which was quantified by a TECAN Spark 10 M microplate reader (TECAN, Switzerland) at emission of 545 nm and excitation of 270 nm. For MHP-induced spore DPA release, the supernatants of spore suspensions after MHP treatment for different times were obtained by centrifugation at 10,000 rpm for 10 min under 4°C. 198 μL of supernatants were mixed with 2 μL of 50 mM TbCl_3_ in a 96-well plate and the fluorescence intensity was immediately measured. For total DPA content, spores (OD_600_ = 0.5) were boiled for 20 min and detected as above [26]. Meanwhile, the DPA standard solution was serially diluted and detected together to obtain a standard curve. The total DPA content was calculated based standard curve.

### Potassium ion release quantification

Potassium ion released by spores under MHP was measured using an Agilent 7800 ICP-MS (Agilent Technologies, Inc., USA) by Shiyanjia Lab (Beijing, China). Specifically, a 2.5 mL aliquot of purified spore suspension (OD_600_ = 2) was sealed in flexible plastic bag, and then subjected to HHP treatment under 200 MPa at 26°C for 1 min. For quantification of total potassium ions in spores, the spore suspensions were lysed by FastPrep-24 (MP Biomedicals, LLC, USA). The supernatants from the pressure-treated and lysed spore suspensions were collected by centrifugation at 12,000 rpm/10 min/4°C, and then filtered through 0.2 μm syringe filters. 1 mL of the supernatant was pipetted into a polytetrafluoroethylene beaker containing 10 mL of 68% nitric acid and heated on a 200°C electric heating plate until organic matter was fully digested. When 1 mL of digestion solution remained, it was diluted with ddH_2_O to a final volume of 10 mL. The concentration of potassium was determined based on standard curve (TMRM^®^ Co., Ltd., Beijing, China). The potassium release percentage was represented as the ratio of released potassium concentration to total potassium concentration.

### Phase-contrast microscopy

Phase-contrast microscopy was performed using a Nikon DS-Qi2 microscope equipped with a Nikon Plan Apo Lambda 100x/1.45 Oil Microscope Objective. Spores (20 μL) were centrifuged, and the pellets were resuspended with 5∼10 μL ddH_2_O and then imaged. Images were analyzed and processed by ImageJ2.

### Modelling of DPA release curves

DPA release curves expressed as %DPA release were fit to BoxLucas1 model using GraphPad Prism 9.5.0 (GraphPad Software Inc., San Diego, CA, USA). The function was expressed as follow:

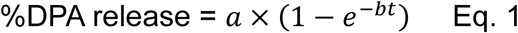

where *t* was HHP treatment time, and *a*, *b* were constants. The germination rate (*K*) at different time were obtained by calculating the derivative of the fitted curves. The product of *a* and *b* was considered as the maximum germination rate (*K*_max_) in the release process. Unless otherwise stated, the maximum germination efficiency (G_max_) was calculated by setting *t* = 10 min in Eq. 1. When germination efficiency was less than 15% after 10-min MHP treatment, G_max_ was indicated as the maximum percentage of DPA release from the corresponding experiment. The adjusted R-square (Adj. R^2^) and root mean square error (RMSE) were calculated to accessed goodness of fit.

### Data processing

Each experiment was performed at least triplicate and the values were represented as mean ± SD. GraphPad Prism 9.5.0 software and IBM SPSS Statistics 26 (SPSS Inc., Ver. 26, Chicago, USA) software were used for statistical analysis, data processing, and graph drawing. One-way ANOVA with Tukey’s post hoc or the independent samples *t* test was performed to compare the significant differences. The significance was established at \**p* < 0.05.

## Acknowledgements

We thank Prof. David Rudner (Harvard Medical School, USA) for generously gifting *B. subtilis* Δ5 and derivative strains. This work was supported by National Natural Science Foundation of China (NSFC) (grant No. 32372470), Agricultural Research Outstanding Talents of China (grant No. 13210317), and 2115 Talent Development Program of China Agricultural University.

## Supplementary Figure Legends

**Figure S1.**
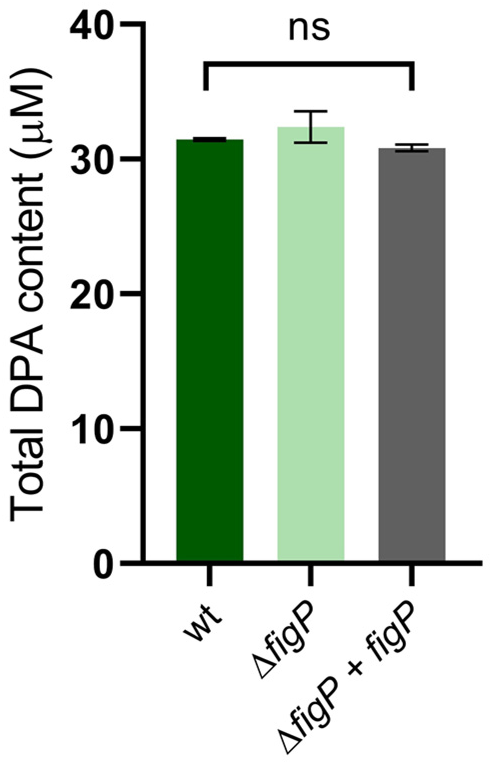
Total DPA content of *B. subtilis* PY79 (wt), YZ03 (Δ*figP*), and YZ05 (Δ*figP, amyE::figP*) spores. Spores (OD=0.5) were boiled for 20 min to determine total DPA content by monitoring the relative fluorescence units (RFU) of Tb^3+^-DPA and then calculating the content based DPA standard curve. Data are represented as mean ± SD. Shown are the representative results of three independent biological experiments, each with three replicates.

**Figure S2.**
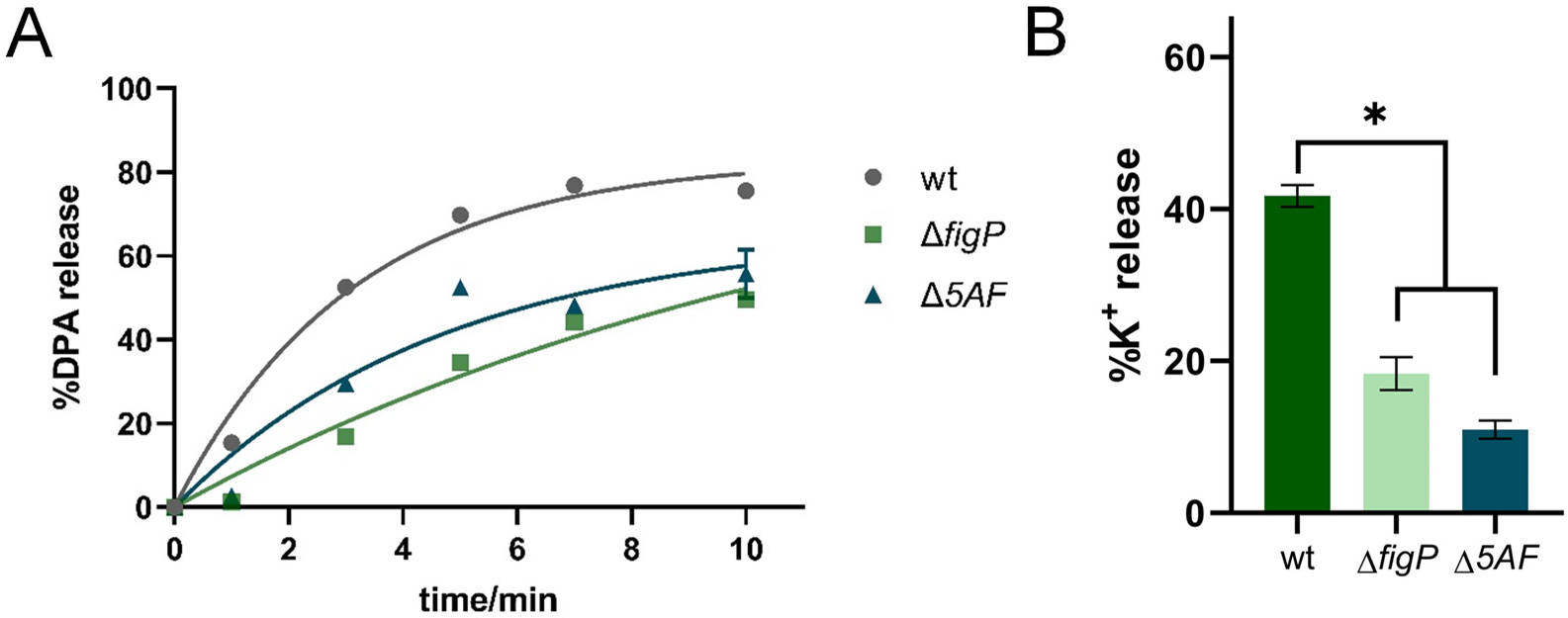
Effects of the absence of SpoVAF or FigP on spore germination at MHP (200 MPa). (A) The percentage DPA release at 0, 1, 3, 5, 7, 10-min MHP treatment, relative to the total DPA content in *B. subtilis* PY79 (wt), YZ03 (Δ*figP*), and KP09 (Δ*spoVAF*) spores. (B) The percentage K^+^ release at 1-min MHP treatment, relative to the total K^+^ content in spores described in (A). Data are represented as mean ± SD. Shown are the representative results of three independent biological experiments, each with three replicates. One-way ANOVA with Tukey’s multiple comparisons test was performed to compare the significant differences. Asterisks denote the significance levels: *p < 0.05.

**Figure S3.**
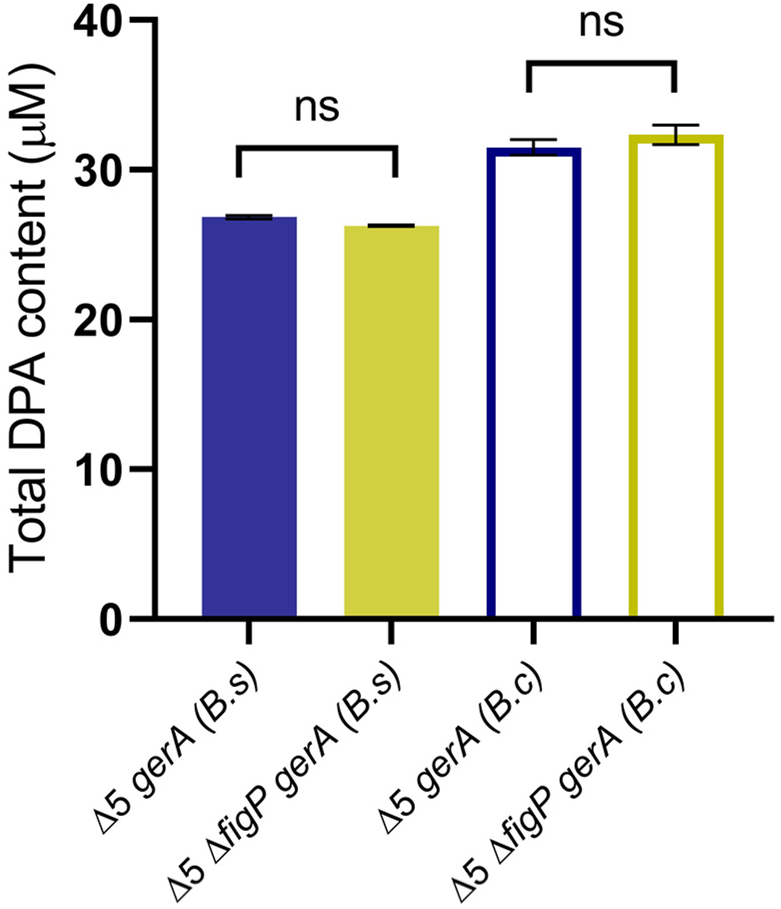
Total DPA content of TYZ12 (Δ5, *amyE::gerA from B. subtilis*), YZR34 (Δ5 Δ*figP*, *amyE::gerA from B. subtilis*), TYZ13 (Δ5, *amyE::gerA from B. cereus*), and YZR37 (Δ5 Δ*figP*, *amyE::gerA from B. cereus*) spores. Spores (OD=0.5) were boiled for 20 min to determine total DPA content by monitoring the relative fluorescence units (RFU) of Tb^3+^-DPA and then calculating the content based DPA standard curve. Data are represented as mean ± SD. Shown are the representative results of three independent biological experiments, each with three replicates.

**Figure S4.**
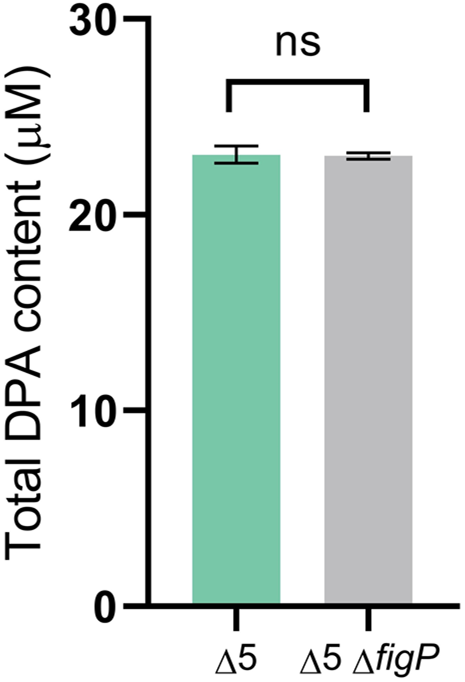
Total DPA content of BLA201 (Δ5) and YZR33 (Δ5 Δ*figP*) spores. Spores (OD=0.5) were boiled for 20 min to determine total DPA content by monitoring the relative fluorescence units (RFU) of Tb^3+^-DPA and then calculating the content based DPA standard curve. Data are represented as mean ± SD. Shown are the representative results of three independent biological experiments, each with three replicates.

**Figure S5.**
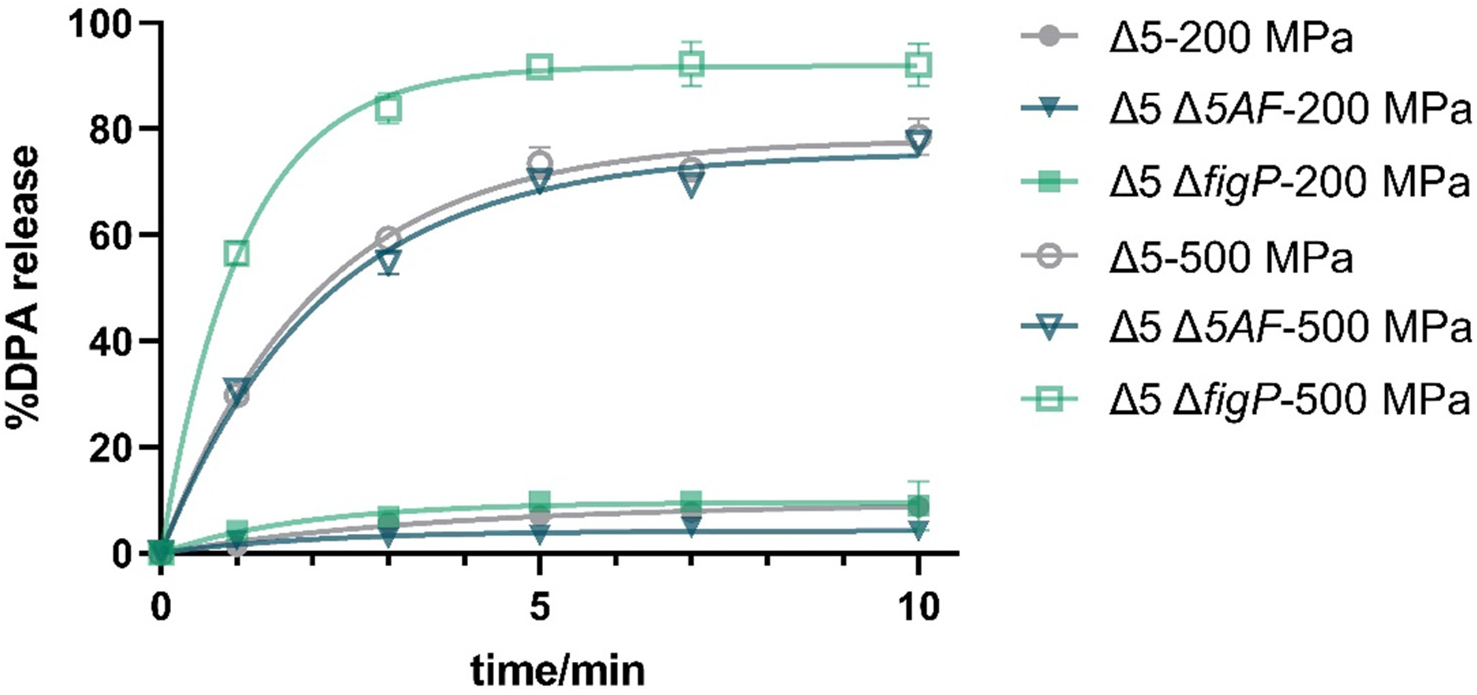
Effects of the absence of SpoVAF or FigP on spore germination at 200 and 500 MPa. Spores of BLA201 (Δ5), YZR33 (Δ5 Δ*figP*) and YZR39 (Δ5 Δ*spoVAF*) strains were treated with 200 and 500 MPa at 26°C to induce germination. DPA release was quantified by monitoring the relative fluorescence units (RFU) of Tb^3+^-DPA, and the percentage of DPA release was calculated as the ratio of the released DPA content to total DPA content. Data are represented as mean ± SD. Shown are the representative results of three independent biological experiments, each with three replicates.

**Figure S6.**
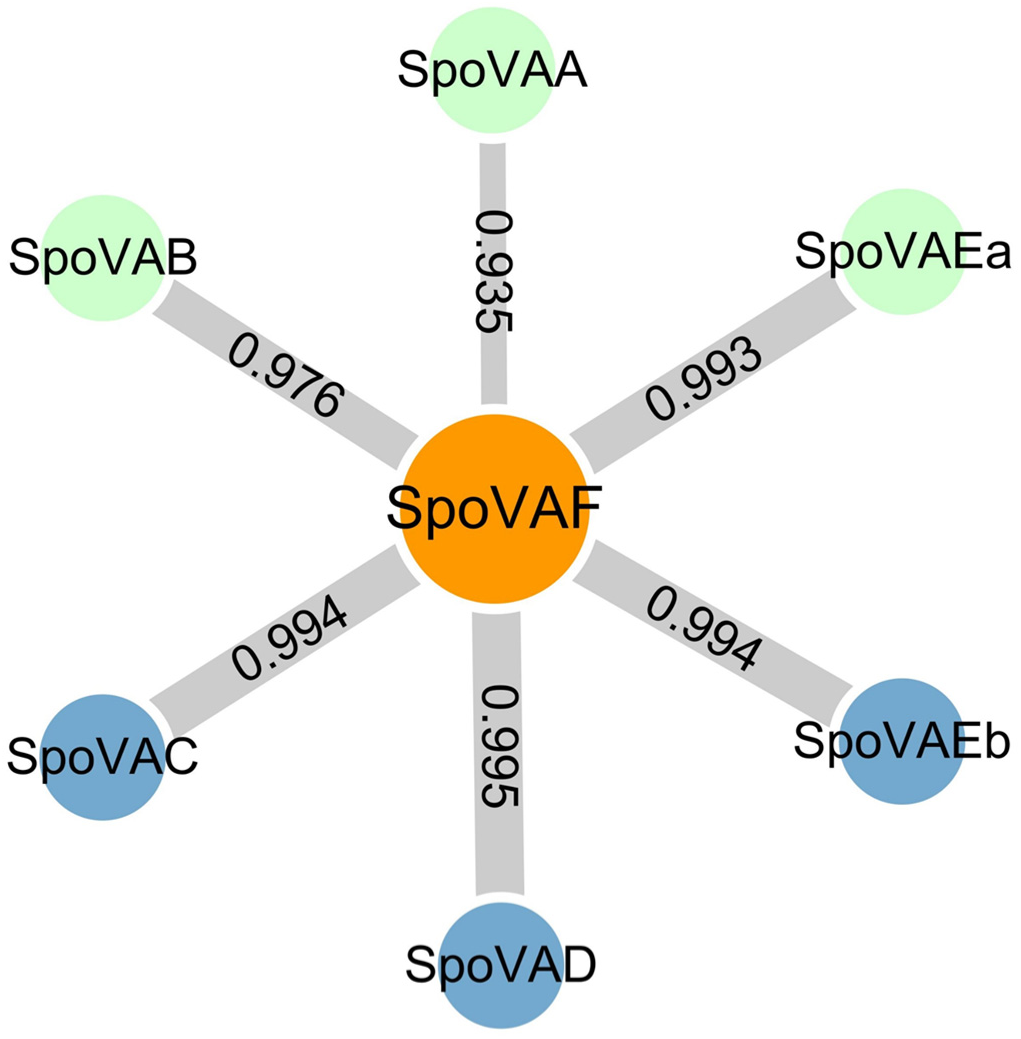
The protein-protein interaction network of SpoVAF and other SpoVA subunits predicted by STRING database. The likelihood of SpoVAF interacting with other subunits was quantified by combined-score, a thicker line indicated a higher prediction score and greater probability of interaction. The SpoVAF subunits (C, D, and Eb) that comprise the DPA release channel were highlighted in blue. SpoVAF exhibited a higher likelihood of interacting with these DPA channel forming subunits.

